# Stability of Oxycodone Solutions Containing S-Ketamine or Dexmedetomidine

**DOI:** 10.64898/2026.04.08.717252

**Authors:** Patrik Väisänen, Sanna Mäkelä, Saija Sirén, Katariina Pohjanoksa, Panu Uusalo, Mika Scheinin, Kirsti Torniainen, Teijo I. Saari

## Abstract

**Objectives:** To determine whether adding *S-*ketamine or dexmedetomidine to oxycodone affects the microbiological, physical, or chemical stability of patient-controlled analgesia (PCA) solutions prepared in a hospital pharmacy.

**Methods:** Oxycodone solution (1 mg/mL) and three oxycodone–*S-*ketamine mixtures (0.25, 0.50, 0.75 mg/mL) and three oxycodone–dexmedetomidine mixtures (2.5, 5.0, 10 µg/mL) were compounded under validated EU GMP Class A/B aseptic conditions and filled into PCA reservoirs. Reservoirs (n=42 for physicochemical studies; n=21 for sterility; n=4 for antimicrobial activity testing) were stored at 2–8°C for 28 days, then at 20–25°C for 2 days. Sterility was assessed by membrane filtration according to Ph. Eur. 2.6.1. Physical stability was evaluated by visual inspection, pH, weight, and osmolality. Chemical stability was assessed using a validated HPLC-UV method developed in accordance with FDA and ICH Q2(R1) guidelines.

**Results:** All antimicrobial activity tests showed growth of the six reference strains, indicating no inhibition by the drug mixtures. All 21 sterility-test reservoirs remained free of turbidity throughout 30 days. No visual changes, precipitation, or discolouration were observed. Weight loss was ≤0.3%, pH changes were between required range 4,5-7, and osmolality increased by <1.4% during the study. Measured oxycodone, *S-*ketamine, and dexmedetomidine concentrations remained within ±5% of initial values, and no degradation products were detected.

**Conclusions:** Oxycodone PCA solutions containing *S-*ketamine or dexmedetomidine remained sterile, physically stable, and chemically stable for 28 days at 2–8°C followed by 2 days at room temperature at 20–25°C. These findings support extended shelf-life and centralized batch preparation of opioid–adjuvant PCA reservoirs in hospital pharmacy practice.

**Key Messages:** *What is already known on this topic:* Opioid–adjuvant combinations such as oxycodone with *S-*ketamine or dexmedetomidine are increasingly used in patient-controlled analgesia, but no commercial multi-agent formulations exist. Hospital pharmacies therefore prepare these mixtures as compounded sterile preparations, despite limited data on their chemical and microbiological stability.

*What this study adds:* This study demonstrates that oxycodone PCA solutions containing *S-*ketamine or dexmedetomidine remain chemically stable and microbiologically sterile for 28 days at 2–8°C plus 2 days at 20–25°C, when prepared under validated aseptic conditions. Concentrations of all analytes remained within ±5% of initial values, and no degradation products were detected using a validated HPLC-UV method.

*How this study might affect research, practice or policy:* These stability data support the assignment of extended beyond-use dates and enable centralized batch compounding of PCA reservoirs in hospital pharmacies. The findings have the potential to reduce aseptic workload, improve production efficiency and decrease medication waste while ensuring product quality.

## 1. INTRODUCTION

Opioids have long been the cornerstone of postoperative pain management but concerns over adverse effects and opioid-related harms have driven increasing interest in multimodal analgesia strategies [1–3]. Patient-controlled analgesia (PCA) enhances pain management by allowing patients to self-administer medication on demand, which often improves patient satisfaction and analgesic control [4,5]. In clinical practice, adjuvant agents such as *S-*ketamine and dexmedetomidine are increasingly combined with opioids in PCA to improve analgesia and reduce opioid requirements [6–10].

However, opioid formulations containing adjuvant drugs are not commercially available. Hospital pharmacies therefore must prepare these mixtures as compounded sterile preparations (CSPs), a process that is technically demanding and subject to strict quality standards [11,12]. For such preparations, reliable data on chemical stability, chemical integrity and microbiological sterility are essential to assign appropriate shelf-lives, ensure patient safety and support efficient production workflows.

Despite the growing clinical use of opioid–adjuvant PCA mixtures, systematic stability data for combinations of oxycodone with *S-*ketamine or with dexmedetomidine remain limited. For hospital pharmacies, this lack of evidence restricts the ability to conduct batch preparation, optimize aseptic workflow, and minimise medication waste. Stability and sterility studies specific to these clinically relevant combinations are therefore needed to guide safe compounding practices and storage conditions.

The aim of the present study was to evaluate the chemical, physical and microbiological stability of oxycodone PCA solutions combined with *S-*ketamine or dexmedetomidine when prepared under validated aseptic conditions and stored in PCA reservoirs under clinically relevant conditions. For this purpose, we first developed and validated a method for determination of concentrations of oxycodone together with *S-*ketamine or dexmedetomidine. After this, we performed a shelf-life study with different combinations of these three drugs used in pain treatment.

## 2. MATERIALS AND METHODS

### 2.2.1 Sample preparation

Description of the drugs, reference solutions and other materials are given in Supplementary text. Oxycodone solution (1 mg/mL as oxycodone hydrochloride trihydrate) was prepared by diluting 10 mL of the pharmaceutical product (Oxanest® 10 mg/mL) into a syringe containing 90 mL of 9 mg/mL (0.9%) sodium chloride solution. Three different mixtures of oxycodone and *S-*ketamine were prepared by mixing known amounts of the pharmaceutical products (Oxanest® 10 mg/mL and Ketanest-S® 25 mg/mL) in syringes containing 9 mg/mL (0.9%) sodium chloride solution (Table 1). Similarly, three different mixtures of oxycodone and dexmedetomidine were prepared by mixing known amounts of the pharmaceutical products (Oxanest® 10 mg/mL and Dexdor® 100 μg/mL) in syringes containing 9 mg/mL (0.9%) sodium chloride solution (Table 1).

**Table 1.** PCA reservoirs containing mixtures of oxycodone, dexmedetomidine and *S-*ketamine.

| **Mixture** | Oxycodone<br>10 mg/mL<br>(mL) | Dexmedetomidine<br>100 µg/mL<br>(mL) | *S*-ketamine<br>25 mg/mL<br>(mL) | 9 mg/mL<br>(0.9%) sodium<br>chloride<br>solution (mL) |
| --- | --- | --- | --- | --- |
| Oxycodone 1 mg/mL +<br>dexmedetomidine 2.5 µg/mL | 10 | 2.5 | - | 87.5 |
| Oxycodone 1 mg/mL +<br>dexmedetomidine 5.0 µg/mL | 10 | 5.0 | - | 85 |
| Oxycodone 1 mg/mL +<br>dexmedetomidine 10.0 µg/mL | 10 | 10.0 | - | 80 |
| Oxycodone 1 mg/mL + *S*-<br>ketamine 0.25 mg/mL | 10 | - | 1 | 89 |
| Oxycodone 1 mg/mL + *S*-ketamine 0.5 mg/mL | 10 | - | 2 | 88 |
| Oxycodone 1 mg/mL + *S*-ketamine 0.75 mg/mL | 10 | - | 3 | 87 |

### 2.2.2 Sample storage

PCA reservoirs were stored for up to 28 days at 2–8 °C in a refrigerator approved for hospital use. Thereafter, they were transferred to room temperature (20–25 °C) for 2 days. The refrigerator was equipped with an automated temperature monitoring system. The PCA reservoirs were protected from light.

Stability of the drug solutions was investigated by analysing samples taken after different periods of storage, on seven different days (days 0, 3, 7, 14, 21, 28 and 30 of the stability test). The study samples consisted of two sets of samples. In the first set, the samples contained oxycodone and *S-*ketamine, and in the second set, they contained oxycodone and dexmedetomidine.

### 2.2.3 Microbiological analysis

Absence of antimicrobial activity of the drug solutions was confirmed with the test solutions containing the highest drug concentrations. Two PCA reservoirs containing oxycodone 1 mg/mL + dexmedetomidine 10 μg/mL and two reservoirs containing oxycodone 1 mg/mL + *S-*ketamine 0.75 mg/mL were filled for this evaluation. The technique of membrane filtration was used as described in Ph. Eur. 2.6.1 [13]. All the samples were withdrawn from PCA reservoirs with a sterile Luer Lock single use syringe. In this procedure, at the beginning the membrane is rinsed with small volume of the sterile solution, followed with the filtration of the sample to be utilized in method suitability test. After this, three repeated rinsings of the membrane are done with the sterile solution. Into each of these last rinsings, a small volume of each microbial strain, to be tested, is added. This test was carried out with six microbial strains, which were: *Aspergillus brasiliensis* (ATCC 16404), *Basillus spizizenii* (ATCC 6633), *Candida albicans* (ATCC 10231), *Clostridium sporogenes* (ATCC 19404), *Pseudomonas aeruginosa* (ATCC 9027) and *Staphylococcus aureus* (ATCC 6538). The filtered volume was 20 mL per membrane. The growth of anaerobic bacteria was examined with membranes incubated in a thioglycolate medium at 30–35 °C. The growth of aerobic bacteria and fungi was examined with membranes incubated in a soybean casein digest medium at 20–25 °C. After 5 days of incubation, the turbidity of each separate media containing six different microbial strains, mentioned above, was observed, according to the requirement of Ph.Eur. Method Suitability Test.

Testing for sterility was carried out using the technique of membrane filtration as described in Ph. Eur. 2.6.1 [13]. For the sterility testing, sample size was 20 ml both for thioglycolate medium and for soybean casein digest medium. All the samples were withdrawn from PCA reservoirs with a sterile Luer Lock single use syringe. The growth of anaerobic bacteria was examined with membranes incubated in a thioglycolate medium at 30–35 °C. The growth of aerobic bacteria and fungi was examined with membranes incubated in a soybean casein digest medium at 20–25 °C. After 14 days of incubation, the turbidity of the media was observed. Twenty-one PCA reservoirs were filled for this assessment, and three parallel PCA reservoirs of each mixture or solution were examined. Sterility samples were aseptically sampled on days 0, 14, 28 and 30. Samples for all microbiological studies were taken and analysed at MetropoliLab (Helsinki, Finland).

### 2.2.4 Physical stability studies

A total of 42 PCA reservoirs were filled to conduct physicochemical evaluations, with six parallel PCA reservoirs for each drug mixture or oxycodone solution. Sample weight and pH were recorded on days 0, 3, 7, 14, 21, 28 and 30. Each PCA reservoir was weighed before and after each sampling day to indicate the possible loss of solvent. A 5 mL sample was taken for pH measurement with a calibrated pH meter (Mettler Toledo SevenCompact S220, Greifensee, Switzerland). The pH meter was calibrated before each test with commercially available buffer solutions. Visual control of each PCA reservoir was performed by observing the samples against a brightly illuminated standard background to determine physical changes in appearance, such as changes in colour, opalescence, precipitation or gas bubble formation.

Osmolality was measured with an automatic osmometer Automatic Osmometer Model 3900 (Advanced Instruments, Norwood, MA, USA). The osmometer was calibrated before each test with a commercially available solution. Osmolality evaluation was conducted on days 0 and 30 at the hospital laboratory of Turku University Hospital. A 2 mL sample was used for osmolality determination. A summary of test schedules is described in Supplementary Table 1.

### 2.2.5 Chemical stability studies

To measure the study samples, we validated an analytical method for the simultaneous determination of oxycodone, *S-*ketamine and dexmedetomidine with reversed-phase high-performance liquid chromatography with ultraviolet detection (HPLC-UV). The analytical method was validated according to FDA [14] and ICH Q2(R1) [15] guidelines; full validation data are provided in the Supplementary material. Validation demonstrated acceptable selectivity, carry-over, linearity, accuracy, precision, and short-term stability across the relevant concentration ranges for oxycodone, *S-*ketamine, and dexmedetomidine.

Quantitation of the analytes was assessed with reversed-phase HPLC-UV. In the validated method no internal standard was used. A sample batch consisted of a blank sample [9 mg/mL (0.9%) sodium chloride solution], eight calibration standards, six QC samples and a set of study samples. The target concentrations in sample analysis were 1000 μg/mL for oxycodone, 250 μg/mL, 500 μg/mL and 750 μg/mL for *S-*ketamine and 2.5 μg/mL, 5 μg/mL and 10 μg/mL for dexmedetomidine.

The quality requirements of the PCA reservoirs in terms of physicochemical and microbiological stability and the test methods used are presented in Table 2.

**Table 2.**
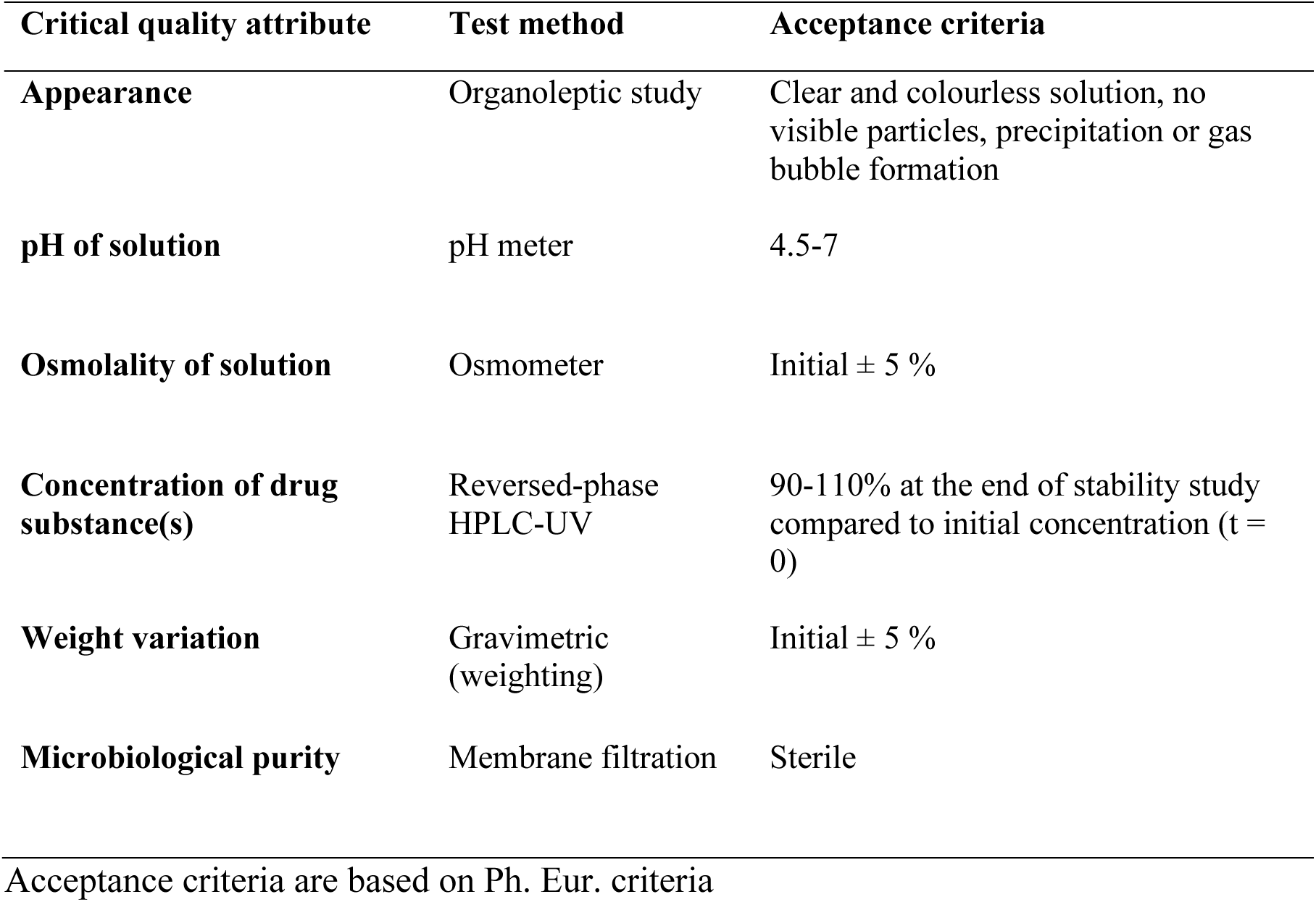
Quality requirements of PCA reservoirs for the stability study.

## 3. RESULTS

### 3.1 Microbiological quality

Absence of antimicrobial activity was demonstrated for all tested drug mixtures (oxycodone 1 mg/mL + dexmedetomidine 10 μg/mL, n = 2 and oxycodone 1 mg/mL + *S-*ketamine 0.75 mg/mL, n = 2) against all of the six reference strains of microbes. The test solutions did not inhibit microbial growth. The results of this microbiological validation indicate that the tested drug mixtures did not interfere with the sterility test results. Thus, the results of the sterility tests can be considered reliable.

All microbiological samples from the 21 PCA reservoirs used for sterility testing were observed to be clear of turbidity over the entire study period. Thus, the studied drug solutions could be considered free of microbial growth and thus sterile. The results indicate, that solutions prepared in EU GMP class B laboratory in class A laminar flow hood of Turku University Hospital Pharmacy, containing oxycodone with or without *S-*ketamine or dexmedetomidine diluted in 9 mg/mL (0.9%) sodium chloride solution, remain sterile and free of microbial growth in PCA reservoirs when stored at 2–8 °C for at least 28 days and thereafter at room temperature (20–25°C) for 2 days, in Turku University Hospital.

### 3.2 Physical stability

All drug solutions were initially clear and colourless and remained unchanged over the study period. No physical changes of appearance, such as visible particles, discolouration, opalescence, precipitation or gas bubble formation were observed in any test sample. Some loss of weight, up to 0.3% per month, was observed in the drug solutions when stored at 2–8 °C for 28 days and thereafter at room temperature (20–25 °C) for 2 days (Table 3). Loss of solvent vehicle was accelerated with increasing temperature. The small obtained loss of weight did not affect the quality and the patient safety of concentrations of the studied substances. The evaporation of solvent was thus considered insignificant.

**Table 3.** Mean weights of the PCA reservoirs.

| Drug solution or mixture | Mean weight (g) | Weight loss during the study (%)<sup>1</sup> |  |  |
| --- | --- | --- | --- | --- |
|  |  | 2-8 °C | 20-25 °C | Total |
| Oxycodone 1 mg/mL | 1635 (1.96) | 0.17 | 0.12 | 0.29 |
| Oxycodone 1 mg/mL +<br>dexmedetomidine 2.5<br>µg/mL | 163 (0.51) | 0.10 | 0.07 | 0.17 |
| Oxycodone 1 mg/mL +<br>dexmedetomidine 5 µg/mL | 164 (0.62) | 0.19 | 0.09 | 0.28 |
| Oxycodone 1 mg/mL +<br>dexmedetomidine 10<br>µg/mL | 165 (0.36) | 0.17 | 0.10 | 0.27 |
| Oxycodone 1 mg/mL +<br>*S*-ketamine 0.25 mg/mL | 165 (0.15) | 0.21 | 0.08 | 0.29 |
| Oxycodone 1 mg/mL +<br>*S*-ketamine 0.50 mg/mL | 163 (0.45) | 0.15 | 0.06 | 0.21 |
| Oxycodone 1 mg/mL +<br>*S*-ketamine 0.75 mg/mL | 164 (0.34) | 0.06 | 0.11 | 0.17 |

<sup>1</sup>Samples were first stored in 2-8 °C for 28 days, after which the reservoirs were transferred to 20-25 °C for an additional two days.
Results are shown as mean (SD). n = 6

The pH of oxycodone solution slightly increased but storage at room temperature (20–25 °C) did not show a significant effect on pH. (Table 4). The pH of mixtures of oxycodone and dexmedetomidine slightly decreased when PCA reservoirs were stored at 2–8°C and then transferred to room temperature (20–25°C). The pH of mixtures of oxycodone and *S-*ketamine slightly increased when PCA reservoirs were stored at 2–8°C. Storage at room temperature (20–25°C) decreased the pH, which was found to be lower than the initial pH. There were no clinically meaningful differences in the pH values between the samples. During the study period, all pH variations were between required range 4.5-7. The measured pH of the PCA reservoirs was always lower than the pKa values of the active substances in them.

**Table 4.** Mean pH values of PCA reservoirs.

| Drug solution or mixture | Initial<sup>1</sup> | 2–8°C | 20–25°C |
| --- | --- | --- | --- |
| Oxycodone 1 mg/mL | 5.61 (0.09)<sup>2</sup> | 5.87 (0.18) | 5.86 (0.05) |
| Oxycodone 1 mg/mL + dexmedetomidine 2.5 µg/mL | 6.11 (0.08) | 5.94 (0.06) | 5.84 (0.03) |
| Oxycodone 1 mg/mL + dexmedetomidine 5 µg/mL | 6.05 (0.04) | 5.92 (0.09) | 5.81 (0.05) |
| Oxycodone 1 mg/mL + dexmedetomidine 10 µg/mL | 6.00 (0.05) | 5.94 (0.09) | 5.89 (0.05) |
| Oxycodone 1 mg/mL + *S*-ketamine 0.25 mg/mL | 5.34 (0.11) | 5.44 (0.06) | 5.22 (0.02) |
| Oxycodone 1 mg/mL + *S*-ketamine 0.50 mg/mL | 5.11 (0.08) | 5.23 (0.06) | 5.00 (0.02) |
| Oxycodone 1 mg/mL + *S*-ketamine 0.75 mg/mL | 5.04 (0.01) | 5.10 (0.08) | 4.90 (0.05) |

<sup>1</sup>Initial refers to day 0 of the study; 2–8°C to day 28, and 20–25°C to day 30 after additional 2 days at room temperature.
<sup>2</sup>Results are presented as mean (SD). n = 6

The osmolality values of all test solutions slightly increased from the initial state during the study period. The increase in osmolality values in the PCA reservoirs was always less than 1.4%, which can be considered insignificant in terms of physical stability. The measured osmolality values were slightly below the conventional reference range of osmolality of human blood plasma, but the differences between the osmolality values measured in the samples were not clinically meaningful.

### 3.3 Chemical stability studies

The results of the analytical method validation proved that our method is appropriate for the determination of concentrations of oxycodone, *S-*ketamine and dexmedetomidine in PCA drug solutions prepared in sterile 9 mg/mL (0.9%) sodium chloride solution. The assay showed linear responses in the tested concentration ranges of 600 – 1200 µg/mL for oxycodone, 200 – 1000 µg/mL for *S-*ketamine and 2 – 12 µg/mL dexmedetomidine. For validation results, see parts 2, 3 and 4 in Supplementary text and Supplementary Tables 2 and 3. Example chromatograms are shown in Figure 1.

**Figure 1.**
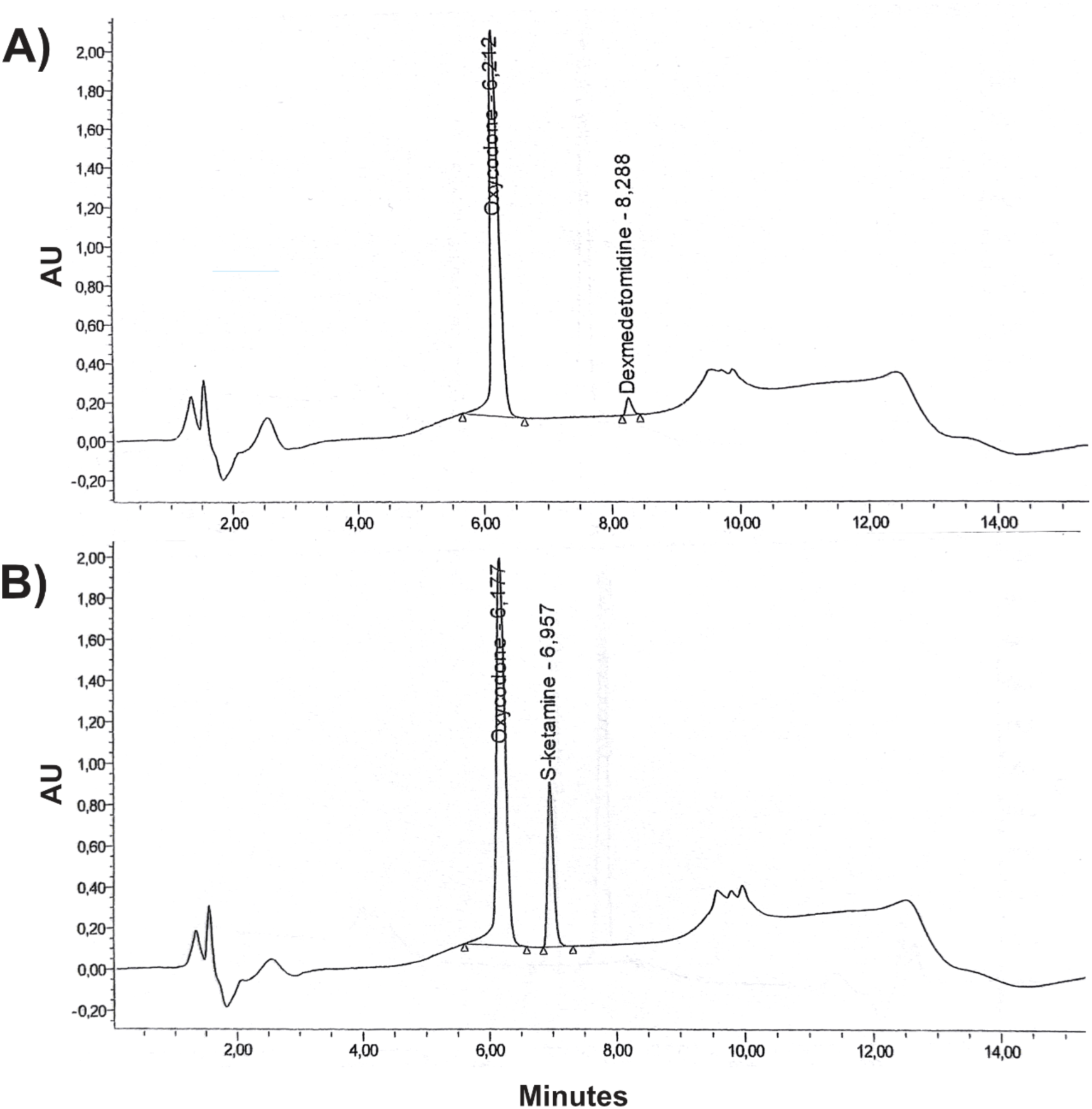
Chromatograms of two quality control samples containing A) oxycodone 1 mg/mL and *S*-ketamine 0.5 mg/mL and B) oxycodone 1 mg/mL and dexmedetomidine 5.0 µg/mL. AU, absorbance unit.

Oxycodone, *S-*ketamine and dexmedetomidine were found to be stable in the vehicle at room temperature at 20–25°C in the laboratory for at least 48 hours and in the autosampler (+20°C) for 24 hours. Mean accuracies of six stability samples were between 95% and 105% compared with the original concentrations.

The chemical stability results are summarised in Table 5, and complete concentration data for all sampling days are presented in Supplementary Tables 4 and 5. Relative concentrations of oxycodone, *S-*ketamine and dexmedetomidine remained within ±5% of the initial concentrations throughout the study period in all PCA reservoirs. These results fulfil the predefined acceptance criteria for chemical stability (Table 2).

**Table 5.** Relative concentrations of oxycodone, *S-*ketamine and dexmedetomidine in PCA reservoirs after 30 days of storage.

| Mixture | Analyte | Relative concentration (% of day 0 result) |
| --- | --- | --- |
|  |  | Mean (SD) |
| Oxycodone 1 mg/mL | Oxycodone | 101.15 (0.40) |
| Oxycodone 1 mg/mL + *S*-ketamine 0.25 mg/mL | Oxycodone | 102.01 (1.27) |
|  | *S*-ketamine | 101.53 (0.50) |
| Oxycodone 1 mg/mL + *S*-ketamine 0.50 mg/mL | Oxycodone | 101.58 (0.41) |
|  | *S*-ketamine | 99.59 (0.44) |
| Oxycodone 1 mg/mL + *S*-ketamine 0.75 mg/mL | Oxycodone | 100.18 (0.31) |
|  | *S*-ketamine | 95.80 (0.48) |
| Oxycodone 1 mg/mL + dexmedetomidine 2.5 µg/mL | Oxycodone | 102.02 (0.63) |
|  | Dexmedetomidine | 96.51 (0.95) |
| Oxycodone 1 mg/mL + dexmedetomidine 5 µg/mL | Oxycodone | 104.08 (0.73) |
|  | Dexmedetomidine | 98.72 (1.22) |
| Oxycodone 1 mg/mL + dexmedetomidine 10 µg/mL | Oxycodone | 104.64 (0.66) |
|  | Dexmedetomidine | 101.42 (1.10) |

## 4. DISCUSSION

The specific aims defined for this stability study were achieved. We established that the employed aseptic working method for the preparation of PCA drug solutions was suitable. Microbiological stability of a PCA solution prepared by a hospital pharmacy can be achieved by following the principles defined in the EU GMP guidelines. An HPLC method for the chemical stability testing of oxycodone, *S-*ketamine and dexmedetomidine was successfully developed.

The HPLC method was suitable and sufficiently reliable for quantitative concentration analysis of oxycodone and *S-*ketamine or dexmedetomidine in a mixture. Oxycodone was found to be physically compatible and chemically stable in mixtures with *S-*ketamine and dexmedetomidine in PCA reservoirs prepared for clinical use [16].

The absence of antimicrobial activity of oxycodone solutions diluted (1 mg/mL) in 9 mg/mL (0.9%) sodium chloride solution has been previously documented [17]. The results of our microbiological studies support the conclusion that oxycodone, *S-*ketamine and dexmedetomidine had no antimicrobial activity which could have interfered with the sterility testing. The results of the microbiological validation indicate that the drug mixtures, at the investigated concentrations, did not interfere with the sterility test results. Thus, the results of the sterility tests can be considered reliable.

Based on the results obtained, the expiration date of the studied drug solution and mixtures in PCA reservoirs can be extended to 28 days when stored in a refrigerator (2–8 °C). Thereafter, the PCA reservoirs can be used by patients for clinical pain management for 2 days. Some loss of weight, up to 0.3% per month, was observed in the drug solution mixtures during the study period. Loss of solvent vehicle occurred during storage, and evaporative loss increased at higher temperatures, but this did not have clinically meaningful effects on the concentrations of the studied substances.

The extent of solvent evaporation was considered insignificant. Similar results of evaporation of solvent have been reported previously in stability studies of PCA reservoirs. A previous study reported loss of solvent at up to 1% per month for fentanyl and sufentanil diluted in 9 mg/mL (0.9%) sodium chloride solution in portable infusion pumps when stored at 4 °C or at room temperature (25 °C) [18]. Amri et al. [17] reported some loss of weight, up to 0.6% per month, for 1 mg/mL oxycodone solutions diluted in 9 mg/mL (0.9%) sodium chloride solution in portable infusion pumps. In their study, different types of infusion pumps were used (CADD and Rythmic reservoirs), and they were stored at room temperature (15–25 °C). Regardless of storage conditions, no visible particles, discolouration, opalescence, precipitation or gas bubble formation were observed in either study.

Storage conditions had no significant impact on the chemical stability of the contents of the PCA reservoirs. According to the physical stability results, the evaporation of solvent during the study period may be related to the observed increase of drug concentrations. No degradation products or impurities were found during the physical stability study period. Possible impurities might be those described in the European Pharmacopoeia for oxycodone (specified impurities A, B, C, D, E and F) and for *S-*ketamine (specified impurities A, B, C and D) [19]. Possible impurity for dexmedetomidine might be levomedetomidine.

Similar results of the chemical stability of oxycodone diluted in 9 mg/mL (0.9%) sodium chloride solution have been reported earlier. Turnbull et al. [20] reported that 5 and 50 mg/mL oxycodone hydrochloride in 9 mg/mL (0.9%) sodium chloride solution retained more than 95% of the initial drug concentrations during 35 days of storage in PVC minibags at 4°C and 35°C.

Only small changes in oxycodone concentrations were also previously reported during 28-day storage of 1 mg/mL oxycodone in 9 mg/mL (0.9%) sodium chloride solution in portable infusion pumps [17]. In that study, two different types of infusion pumps were (CADD and Rythmic^TM^ reservoirs) kept at room temperature (15–25°C). Oxycodone concentrations decreased 1.4% in CADD reservoirs and increased 1.5% in Rythmic^TM^ reservoirs [17]. In the same study, oxycodone concentrations decreased 3.9% in Rythmic^TM^ reservoirs protected from light, but no impact on product stability was noted. Because of the use of different infusion pumps, the previous results are not fully comparable with our setting. For example, evaporation of water may vary between different infusion pumps and their reservoirs.

The stability of oxycodone solutions in syringes used for long-duration infusions has been evaluated earlier [17, 20]. A previous report stated that oxycodone hydrochloride prepared in sterile water retained more than 95% of the original concentration during 35 days of storage at 4° C and 35° C [17]. The same was observed with oxycodone solutions diluted with different solvents [20]. More relevant to the current study, a long-term stability study of oxycodone solutions stored in a PCA reservoir over 28 days showed no significant changes in pH, weight or osmolality [17]. Similar results have been reported for ketamine and dexmedetomidine. Daouphans et al. [21] reported that *S-*ketamine (0.1–40 mg/mL) combined with oxycodone (0.4–10 mg/mL) was chemically stable for 7 days at room temperature (23°C) when diluted in 9 mg/ml (0.9%) sodium chloride solution and packaged in polypropylene syringes or PVC bags. All formulations maintained more than 95% of their initial drug concentrations over the study period.

Chromatograms showed no detectable degradation products for any analyte. Solutions of ketamine combined with morphine were physicochemically stable at 5° C and 23° C for 91 days when diluted with 9 mg/mL (0.9%) sodium chloride solution [22]. Exposure to light had no impact on the stability of these drugs, as their concentrations remained above 98% of the original concentration under both storage conditions [22]. Mixtures of tramadol with ketamine in 9 mg/mL (0.9%) sodium chloride solution prepared for PCA delivery systems were stable for 14 days when stored at 4°C and 25°C [23].

Stability of four different concentrations of dexmedetomidine hydrochloride (4, 8, 12 and 20 µg/mL) have been tested in PVC bags containing 9 mg/mL (0.9%) sodium chloride solution [24]. Solutions remained stable at room temperature at 20–25°C, over the 48-hour testing period, and only small pH decreases were seen with increasing dexmedetomidine concentrations [24].

These findings have direct implications for hospital pharmacy practice. Demonstrating that oxycodone PCA formulations containing *S-*ketamine or dexmedetomidine remain chemically stable and sterile for 28 days at 2–8°C (plus 2 days at room temperature at 20–25°C) supports the use of extended beyond-use dates and enables centralized batch compounding. This can reduce aseptic workload, improve utilization of cleanroom capacity, and decrease material and drug waste. The validated analytical method and consistent sterility results provide evidence-based assurance of product quality throughout the storage period. As demand for opioid–adjuvant PCA combinations increases, such stability data are essential for safe, efficient, and sustainable provision of compounded sterile preparations in hospital pharmacies.

## DISCLOSURES

### Ethics approval

Not applicable. This study was a laboratory-based investigation of drug solution stability and did not involve human participants, patient data, or animal experiments; therefore, ethical approval was not required.

### Data management and sharing

Original study data may be available to qualified investigators upon reasonable request addressed to the corresponding author.

### AI technology

During the preparation of this work the authors used ChatGPT to improve language and readability. After using this tool/service, the authors reviewed and edited the contents as needed and take full responsibility for the contents of the publication.

## Acknowledgements

The authors wish to acknowledge the late Lauri Vuorilehto, MSc for his significant contribution to this work. The Hospital Pharmacy of Turku University Hospital is acknowledged for assistance, and ward pharmacists Anne Honkanen, Kira Honkanen and Elina Luoto are thanked for fruitful collaboration.

## SUPPLEMENTARY TEXT

### 1. Materials

The supplementary material contains detailed analytical validation data and complete concentration datasets supporting the results presented in the main manuscript.

#### 1.1. Drugs

An oxycodone solution formulation for injection (Oxanest® 10 mg/mL, Takeda Oy, Helsinki, Finland) containing 100 mg of oxycodone hydrochloride trihydrate in 10 mL isotonic solution in water for injection (containing oxycodone at 7.8 mg/mL). *S-*ketamine concentrate for solution (Ketanest-S® 25 mg/mL, Pfizer Oy, Helsinki, Finland) containing 50 mg of *S*-ketamine in 2 mL isotonic solution in water for injection. Dexmedetomidine concentrate for infusion (Dexdor®, 100 µg/mL, Orion Corporation, Espoo, Finland) containing 200 µg of dexmedetomidine in 2 mL isotonic solution in water for injection. All drugs were preservative-free and were acquired from the manufacturers by Turku University Hospital Pharmacy.

For clarity, it should be noted that the concentrations reported for oxycodone refer to oxycodone hydrochloride trihydrate; however, throughout the text, they are referred to simply as oxycodone concentrations for simplicity.

#### 1.2. Reference solutions

Commercially available pharmaceutical products listed above in 1.1 were used as reference solutions in the concentration analysis of oxycodone and dexmedetomidine. For *S*-ketamine Ketanest-S^®^ 5 mg/mL (Pfizer Oy, Helsinki, Finland) was used. Reference solutions were used for the preparation of calibration standards and quality control samples with adequate concentrations for the chemical stability study. NaCl 0.9% infusion solution (Natriumklorid B. Braun 9 mg/mL, B. Braun Melsungen AG, Melsungen, Germany) was used as analyte-free matrix for preparation of calibration standards and quality control samples, and as blank samples to serve as vehicle control samples. Calibration standards and QC samples of the validation procedure contained all three analytes (oxycodone, *S-*ketamine and dexmedetomidine), but the study sample batches contained only the same two analytes that were present in the study samples (oxycodone and *S-*ketamine or oxycodone and dexmedetomidine). The volume of the missing analyte was replaced with 0.9% NaCl solution.

#### 1.3. Other materials

CADD-Legacy^TM^ PCA (Steripolar Oy, Espoo, Finland) medication cassette reservoirs were used as pharmaceutical packaging material. These cassettes consisted of a medical-grade infusion bag made of polyvinyl chloride (PVC) with an inner layer of phthalate ester placed in a light-protecting polycarbonate reservoir, and a Luer-Lock® PVC tube with the plasticizer di-(2-ethylhexyl) phthalate (DEHP). Millipore Millex-GS® (Merck, Darmstadt, Germany) sterile syringe filters were used for sterilization of sample solutions after preparation (membrane filtration). The syringe filters had a pore size of 0.22 μm and consisted of hydrophilic mixed cellulose esters. Compounding and PCA reservoir filling are described in the Supplementary text, part 2.2.

### 2. Validation of an analysis method for determination of concentrations of oxycodone, *S*-ketamine and dexmedetomidine

Method validation for selectivity, carry-over, linearity, precision, accuracy and short-term stability of solutions was performed using dilutions of authentic reference solutions. Method validation was performed according to Guidance on Analytical Procedures and Methods Validation for Drugs and Biologics (FDA 2015) and ICH Q2 (R1) Guidance on Validation of Analytical Procedures: Text and Methodology (ICH 2005). The study was subject to quality assurance evaluation by the Quality Assurance inspector of the test facility. The inspector inspected the Analytical Study Plan and its Amendment No. 1 and the critical points of the technical performance of the study.

#### 2.1. Materials

Reference items (Oxanest 10 mg/mL, Takeda Oy; Ketanest-S 5 mg/mL, Pfizer Oy; Dexdor 100 µg/mL, Orion Oyj) and a 9 mg/ml (0.9 %) sodium chloride (Natriumklorid B. Braun 9 mg/mL, B. Braun Melsungen AG) were used for preparation of calibration standards, quality control and blank samples. Reference solutions, calibration standards and quality control samples were stored at room temperature, protected from light. Reference items were provided by Turku University Hospital.

#### 2.2. Sample preparation

Oxycodone solution (1 mg/mL) was prepared by diluting 10 mL of the pharmaceutical product (Oxanest® 10 mg/mL) into a syringe containing 90 mL of 9 mg/mL NaCl solution. Three different mixtures of oxycodone and *S*-ketamine were prepared by mixing known amounts of the pharmaceutical products (Oxanest® 10 mg/mL and Ketanest-S® 25 mg/mL) in syringes containing 9 mg/mL NaCl solution (Table 1). Similarly, three different mixtures of oxycodone and dexmedetomidine were prepared by mixing known amounts of the pharmaceutical products (Oxanest® 10 mg/mL and Dexdor® 100 μg/mL) in syringes containing 9 mg/mL NaCl solution (Table 1).

After mixing and dilution, the final volume of each sample was 100 ml. The whole contents of the syringes were transferred into PCA reservoirs to investigate their microbiological and physicochemical stability. The transfer of samples into the PCA reservoirs was done through a membrane filter. A filter integrity test was performed with a bubble-point test apparatus for each membrane filter used. All compoundings and PCA reservoir fillings were performed under controlled and validated aseptic conditions in an EU GMP class B room and class A laminar flow hood.

#### 2.3. Drug concentration analysis

Quantitation of the analytes was accomplished with validated reversed-phase HPLC-UV. No internal standard was used. Drug concentration analysis of the study samples was performed on the same 42 PCA reservoirs filled for the physicochemical stability studies. An approximate volume of 1.5 mL of each PCA reservoir was aseptically sampled using a polypropylene syringe and collected into a 1.5 mL Eppendorf tube made of polypropylene on days 0, 3, 7, 14, 21, 28 and 30. At least 100 μL sample volumes were used for the HPLC-UV analysis. The study samples were analysed without any sample preparation.

Chromatographic separations were performed with a SunFire^TM^ analytical column (C18, 3.5 μm, 2.1 x 150 mm, Waters) coupled with a Gemini^®^ guard column (C18, 2 x 4 mm, Phenomenex). The mobile phase consisted of two eluents: A was 0.1% formic acid in water and B was 0.1% formic acid in acetonitrile. A chromatographic run with a gradient was used: from 0 min to 0.2 min A was 97% (isocratic); from 0.2 min to 5 min A was decreased from 97% to 70%, from 5 min to 6.5 min A was decreased from 70% to 10%, from 6.5 min to 8.5 min A was 10% (isocratic), from 8.5 min to 9 min A was increased to 97% and lastly A was held constant at 97% from 9 min to 15.5 min. The flow rate was 300 μL/min resulting in retention times of approximately 6.4 min for oxycodone, 7.1 min for *S*-ketamine and 8.4 min for dexmedetomidine. The temperature of the autosampler sample compartment was set at 20 °C and the column oven temperature was set to 25 °C. The injection volume was 5 μL. A Waters Alliance 2695 separations module connected with a Waters 2487 dual λ Absorbance Detector set at a detection wavelength of 220 nm was used. Each sample was injected twice and the average concentration was reported as the result. Detailed working instructions for preparation of the validation samples and the study samples for HPLC-UV analysis were approved before commencement of the study sample analysis.

### 3. Validation study

Method validation for selectivity, carry-over, linearity, precision, accuracy and short-term stability of solutions was performed using dilutions of authentic reference solutions. Method validation was performed according to relevant regulatory guidance [14, 15].

#### 3.1. Selectivity and carry-over

Potential sources of interference were evaluated to test the selectivity and carry-over effect of the method. In three validation batches analyte-free matrix (blank sample) was analysed prior to calibration standards, after the highest calibration standard and at the end of the batch. According to the results, no interferences or carry-over effects were detected. Moreover, the chromatograms of the stored samples were visually compared with the chromatograms of freshly prepared calibration standard samples to detect possible additional peaks indicating degradation products. No additional chromatographic peaks were observed during the stability study.

#### 3.2. Linearity

The linearity of the method was tested by preparing calibration standard samples ST1–ST8 at eight different concentrations in the target ranges (oxycodone 0.6-1.2 mg/mL; *S*-ketamine 0.2-1.0 mg/mL; dexmedetomidine 2-12 µg/mL). Calibration standards were prepared and analysed three times. Calibration curves were generated using weighted regression analysis, and the adequacy of the curve fitting was assessed with residual plots and correlation coefficients (r^2^).

The validation assay showed linear responses in the concentration ranges of 600–1200 μg/mL for oxycodone, 200–1000 μg/mL for *S-*ketamine and 2–12 μg/mL for dexmedetomidine (Supplementary s 4). The acceptance criteria for the inter-assay relative bias of all back-calculated values of the calibration standards was set between -5% and +5%. The criteria were met.

#### 3.3. Precision and accuracy

Inter- and intra-assay precision and accuracy were evaluated to three quality control samples at three concentration levels of each studied substance. Quality control samples were assigned as low (QC1), medium (QC2) and high concentration QC (QC3) and their oxycodone, *S*-ketamine and dexmedetomidine concentrations were 1.0 mg/mL, 0.25 mg/mL and 2.5 µg/mL for QC1, 1.0 mg/mL, 0.5 mg/mL and 5 µg/mL for QC2 and 1.0 mg/mL, 0.75 mg/mL and 10 µg/mL for QC3, respectively. In total, three validation batches were analysed and in each batch, six replicates of all control samples were measured. The precision was determined as the intra-assay and inter-assay relative standard deviations (RSD) of the determined concentrations in percentages: RSD (%) = (SD/M) × 100, where SD is the standard deviation and M is the mean of the experimentally obtained concentrations.

We report accuracy as follows: Accuracy (%) = mean/nominal concentration*100. The validation data are summarized in Supplementary Tables 2 and 3. In the validation assays, accuracies and precisions of QC samples were always between 95% and 105% with an RSD of < 5% compared to the nominal value.

### 4. Data analysis

The calculations for the quantitation were based on peak areas of the analytes. The data from the HPLC-UV analyses were collected using Waters Empower 3 software. The peak integrations, calibration curves and quantitations were generated with the same software. The standard curves were generated using linear regression with 1/x^2^ weighting.

## Supplementary Tables

**Supplementary Table 1.** A summary of test schedules.

|  | **Day 0** | **Day 3** | **Day 7** | **Day 14** | **Day 21** | **Day 28** | **Day 30** |
| --- | --- | --- | --- | --- | --- | --- | --- |
| **Microbiological quality** |  |  |  |  |  |  |  |
| Sterility | x |  |  | x |  | x | x |
| **Physical characteristics** |  |  |  |  |  |  |  |
| Weight | x | x | x | x | x | x | x |
| pH | x | x | x | x | x | x | x |
| Visual control | x | x | x | x | x | x | x |
| Osmolality | x |  |  |  |  |  | x |
| **Chemical quality** |  |  |  |  |  |  |  |
| Drug concentrations | x | x | x | x | x | x | x |

**Supplementary Table 2.** Accuracy and precision of the back-calculated calibration standards. Each calibration standard was analysed once in three validation batches.*

| Drug | Nominal concentration (µg/mL) |  |  |  |  |  |  |  |
| --- | --- | --- | --- | --- | --- | --- | --- | --- |
| Oxycodone | 600 µg/mL | 700 µg/mL | 800 µg/mL | 900 µg/mL | 1000 µg/mL | 1050 µg/mL | 1100 µg/mL | 1200 µg/mL |
| Mean (SD) | 597.0 (5.0) | 697.6 (6.0) | 805.3 (5.0) | 906.6 (3.9) | 1010.3 (3.1) | 1053.1 (7.7) | 1090.4 (3.2) | 1187.9 (8.7) |
| RSD (%) | 0.830 | 0.864 | 0.618 | 0.434 | 0.306 | 0.731 | 0.290 | 0.731 |
| Bias (%) | -0.49 | -0.34 | 0.66 | 0.74 | 1.03 | 0.29 | -0.87 | -1.01 |
| *S*-ketamine | 200 µg/mL | 235 µg/mL | 300 µg/mL | 450 µg/mL | 600 µg/mL | 750 µg/mL | 900 µg/mL | 1000 µg/mL |
| Mean (SD) | 200.2 (0.1) | 235.0 (0.3) | 298.9 (1.0) | 450.9 (1.6) | 603.8 (0.3) | 746.2 (2.6) | 892.2 (1.6) | 1008.2 (0.8) |
| RSD (%) | 0.029 | 0.130 | 0.335 | 0.351 | 0.053 | 0.342 | 0.184 | 0.077 |
| Bias (%) | 0.08 | 0.01 | -0.36 | 0.19 | 0.64 | -0.51 | -0.87 | 0.82 |
| Dexmedetomidine | 2.0 µg/mL | 2.3 µg/mL | 3.0 µg/mL | 5.0 µg/mL | 7.0 µg/mL | 9.0 µg/mL | 11.0 µg/mL | 12.0 µg/mL |
| Mean (SD) | 2.0 (0.0) | 2.4 (0.0) | 3.0 (0.0) | 5.0 (0.0) | 7.0 (0.0) | 9.0 (0.0) | 11.0 (0.1) | 12.0 (0.1) |
| RSD (%) | 0.338 | 0.534 | 0.451 | 0.412 | 0.382 | 0.197 | 0.820 | 0.625 |
| Bias (%) | -1.52 | 2.36 | -0.78 | -0.82 | 0.38 | 0.23 | 0.08 | -0.19 |

Results are shown as mean, standard deviation (SD), relative standard deviation (RSD) and relative bias.
\*The coefficients of determination ( $r^2$ ) were 0.998–0.999, 1.000 and 0.999–1.000 for oxycodone, *S*-ketamine and dexmedetomidine, respectively.

**Supplementary Table 3.** Validation data for accuracy and precision of the analytical method for the measurement of concentrations of oxycodone, *S*-ketamine and dexmedetomidine in the quality control samples (QC1−QC3).

| Intra-assay |  |  |  |  |
| --- | --- | --- | --- | --- |
| Nominal concentration of quality control sample (µg/mL) | Mean | RSD (%) | Accuracy (%) | n |
| **Oxycodone** |  |  |  |  |
| 1000 µg/mL (QC1) | 996.0 – 997.9 | 0.3 – 0.6 | 99.6 – 99.8 | <sup>a</sup> |
| 1000 µg/mL (QC2) | 993.1 – 998.0 | 0.4 – 0.9 | 99.3 – 99.8 | <sup>a</sup> |
| 1000 µg/mL (QC3) | 993.2 – 1002.5 | 0.5 – 0.8 | 99.3 – 100.3 | <sup>a</sup> |
| ***S*-Ketamine** |  |  |  |  |
| 250 µg/mL (QC1) | 243.4 – 247.5 | 0.5 – 0.8 | 97.4 – 99.0 | <sup>a</sup> |
| 500 µg/mL (QC2) | 492.8 – 496.1 | 0.6 | 98.6 – 99.2 | <sup>a</sup> |
| 750 µg/mL (QC3) | 737.0 – 744.5 | 0.2 – 0.8 | 98.3 – 99.3 | <sup>a</sup> |
| **Dexmedetomidine** |  |  |  |  |
| 2.5 µg/mL (QC1) | 2.5 | 0.8 – 2.8 | 99.6 – 100.6 | <sup>b</sup> |
| 5.0 µg/mL (QC2) | 4.9 – 5.0 | 0.3 – 0.9 | 97.1 – 99.3 | <sup>a</sup> |
| 10 µg/mL (QC3) | 9.8 – 10.0 | 0.4 – 0.8 | 98.4 – 99.5 | <sup>a</sup> |
| Inter-assay |  |  |  |  |
| **Oxycodone** |  |  |  |  |
| 1000 µg/mL (QC1) | 997.2 | 0.5 | 99.7 | 18 |
| 1000 µg/mL (QC2) | 995.5 | 0.6 | 99.5 | 18 |
| 1000 µg/mL (QC3) | 996.4 | 0.7 | 99.6 | 18 |
| ***S*-Ketamine** |  |  |  |  |
| 250 µg/mL (QC1) | 245.4 | 0.9 | 98.2 | 18 |
| 500 µg/mL (QC2) | 494.3 | 0.6 | 98.9 | 18 |
| 750 µg/mL (QC3) | 741.4 | 0.8 | 98.9 | 18 |
| **Dexmedetomidine** |  |  |  |  |
| 2.5 µg/mL (QC1) | 2.5 | 1.6 | 100.1 | 17 |
| 5.0 µg/mL (QC2) | 4.9 | 1.1 | 98.2 | 18 |
| 10 µg/mL (QC3) | 9.9 | 0.7 | 98.8 | 18 |

<sup>a</sup> six replicates per assay, three assays in total
<sup>b</sup> six replicates per assay, three assays in total. One replicate rejected as an outlier.
RSD, relative standard deviation

**Supplementary Table 4.**
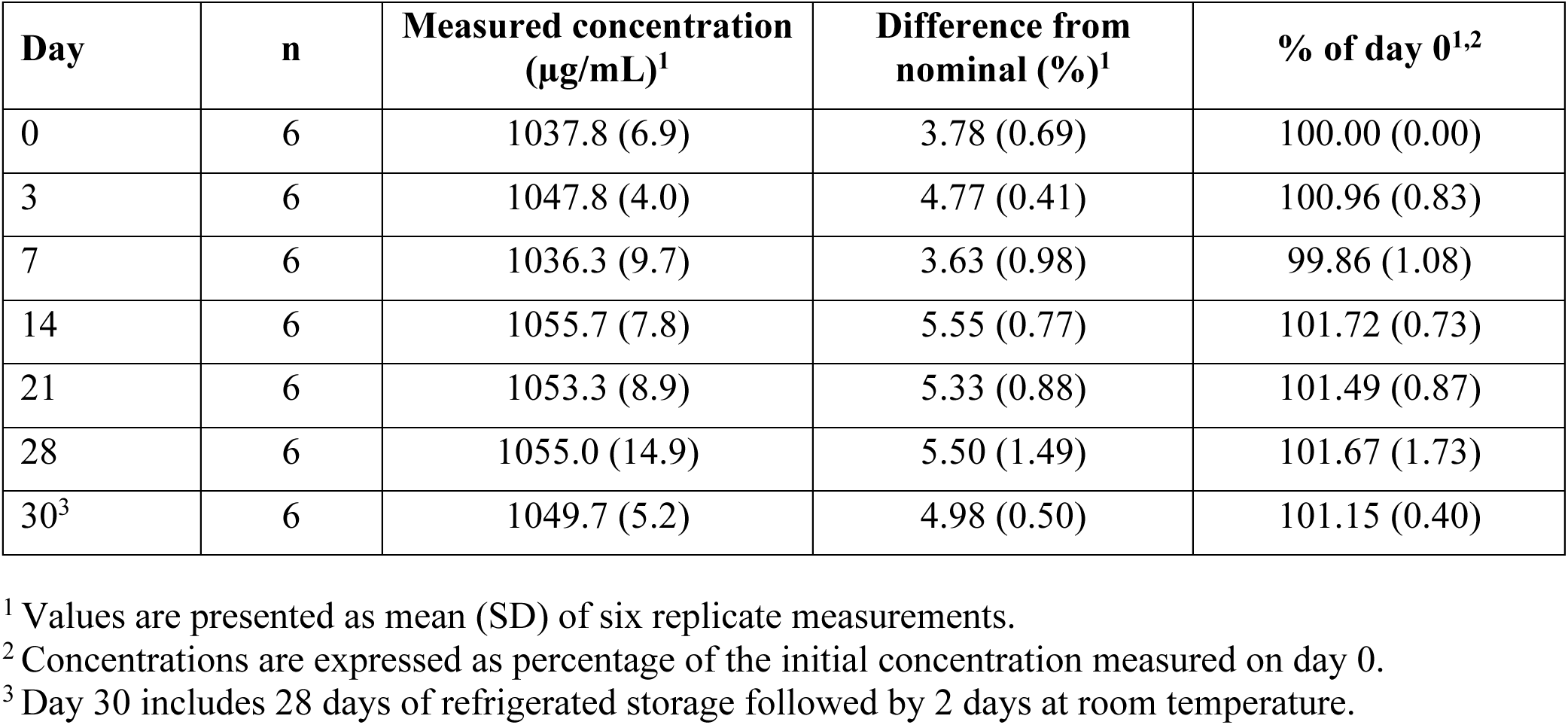
Complete concentration data of oxycodone solution during the stability study.

**Supplementary Table 5.** Complete concentration data of oxycodone mixtures containing *S-*ketamine or dexmedetomidine during the stability study^1^.

| Mixture | Analyte | Day | n | Initial concentration (µg/mL) | Measured concentration (µg/mL) | % of day 0<sup>2</sup> |
| --- | --- | --- | --- | --- | --- | --- |
| Oxycodone 1 mg/mL + *S*-ketamine 0.25 mg/mL | Oxycodone | 0 | 6 | 1000.0 | 1000.0 (0.0) | 100.00 (0.00) |
| Oxycodone 1 mg/mL + *S*-ketamine 0.25 mg/mL | Oxycodone | 3 | 6 | 1000.0 | 999.1 (9.1) | 99.91 (0.91) |
| Oxycodone 1 mg/mL + *S*-ketamine 0.25 mg/mL | Oxycodone | 7 | 6 | 1000.0 | 1004.7 (3.1) | 100.47 (0.31) |
| Oxycodone 1 mg/mL + *S*-ketamine 0.25 mg/mL | Oxycodone | 14 | 6 | 1000.0 | 1005.7 (5.7) | 100.57 (0.57) |
| Oxycodone 1 mg/mL + *S*-ketamine 0.25 mg/mL | Oxycodone | 21 | 6 | 1000.0 | 1001.7 (7.1) | 100.17 (0.71) |
| Oxycodone 1 mg/mL + *S*-ketamine 0.25 mg/mL | Oxycodone | 28 | 6 | 1000.0 | 1020.3 (7.6) | 102.03 (0.76) |
| Oxycodone 1 mg/mL + *S*-ketamine 0.25 mg/mL | Oxycodone | 30<sup>3</sup> | 6 | 1000.0 | 1020.1 (12.7) | 102.01 (1.27) |
| Oxycodone 1 mg/mL + *S*-ketamine 0.25 mg/mL | *S*-Ketamine | 0 | 6 | 250.0 | 250.0 (0.0) | 100.00 (0.00) |
| Oxycodone 1 mg/mL + *S*-ketamine 0.25 mg/mL | *S*-Ketamine | 3 | 6 | 250.0 | 253.8 (1.0) | 101.51 (0.41) |
| Oxycodone 1 mg/mL + *S*-ketamine 0.25 mg/mL | *S*-Ketamine | 7 | 6 | 250.0 | 257.2 (0.9) | 102.87 (0.34) |
| Oxycodone 1 mg/mL + *S*-ketamine 0.25 mg/mL | *S*-Ketamine | 14 | 6 | 250.0 | 254.3 (1.2) | 101.73 (0.48) |
| Oxycodone 1 mg/mL + *S*-ketamine 0.25 mg/mL | *S*-Ketamine | 21 | 6 | 250.0 | 256.8 (0.4) | 102.72 (0.16) |
| Oxycodone 1 mg/mL + *S*-ketamine 0.25 mg/mL | *S*-Ketamine | 28 | 6 | 250.0 | 258.4 (1.1) | 103.35 (0.44) |
| Oxycodone 1 mg/mL + *S*-ketamine 0.25 mg/mL | *S*-Ketamine | 30<sup>3</sup> | 6 | 250.0 | 253.8 (1.3) | 101.53 (0.50) |
| Oxycodone 1 mg/mL + *S*-ketamine 0.50 mg/mL | Oxycodone | 0 | 6 | 1000.0 | 1000.0 (0.0) | 100.00 (0.00) |
| Oxycodone 1 mg/mL + *S*-ketamine 0.50 mg/mL | Oxycodone | 3 | 6 | 1000.0 | 1008.1 (13.3) | 100.81 (1.33) |
| Oxycodone 1 mg/mL + *S*-ketamine 0.50 mg/mL | Oxycodone | 7 | 6 | 1000.0 | 988.7 (5.4) | 98.87 (0.54) |
| Oxycodone 1 mg/mL + *S*-ketamine 0.50 mg/mL | Oxycodone | 14 | 6 | 1000.0 | 1003.9 (6.3) | 100.39 (0.63) |
| Oxycodone 1 mg/mL + *S*-ketamine 0.50 mg/mL | Oxycodone | 21 | 6 | 1000.0 | 1005.1 (8.4) | 100.51 (0.84) |
| Oxycodone 1 mg/mL + *S*-ketamine 0.50 mg/mL | Oxycodone | 28 | 6 | 1000.0 | 1005.8 (7.9) | 100.58 (0.79) |
| Oxycodone 1 mg/mL + *S*-ketamine 0.50 mg/mL | Oxycodone | 30<sup>3</sup> | 6 | 1000.0 | 1015.8 (4.1) | 101.58 (0.41) |
| Oxycodone 1 mg/mL + *S*-ketamine 0.50 mg/mL | *S*-Ketamine | 0 | 6 | 500.0 | 500.0 (0.0) | 100.00 (0.00) |
| Oxycodone 1 mg/mL + *S*-ketamine 0.50 mg/mL | *S*-Ketamine | 3 | 6 | 500.0 | 494.6 (2.6) | 98.92 (0.52) |
| Oxycodone 1 mg/mL + *S*-ketamine 0.50 mg/mL | *S*-Ketamine | 7 | 6 | 500.0 | 498.2 (1.9) | 99.63 (0.37) |
| Oxycodone 1 mg/mL + *S*-ketamine 0.50 mg/mL | *S*-Ketamine | 14 | 6 | 500.0 | 500.1 (2.0) | 100.02 (0.40) |
| Oxycodone 1 mg/mL + *S*-ketamine 0.50 mg/mL | *S*-Ketamine | 21 | 6 | 500.0 | 499.0 (2.9) | 99.80 (0.59) |
| Oxycodone 1 mg/mL + *S*-ketamine 0.50 mg/mL | *S*-Ketamine | 28 | 6 | 500.0 | 496.9 (2.4) | 99.37 (0.48) |
| Oxycodone 1 mg/mL + *S*-ketamine 0.50 mg/mL | *S*-Ketamine | 30<sup>3</sup> | 6 | 500.0 | 497.9 (2.2) | 99.59 (0.44) |
| Oxycodone 1 mg/mL + S-ketamine 0.75 mg/mL | Oxycodone | 0 | 6 | 1000.0 | 1000.0 (0.0) | 100.00 (0.00) |
| Oxycodone 1 mg/mL + S-ketamine 0.75 mg/mL | Oxycodone | 3 | 6 | 1000.0 | 1017.4 (8.1) | 101.74 (0.81) |
| Oxycodone 1 mg/mL + S-ketamine 0.75 mg/mL | Oxycodone | 7 | 6 | 1000.0 | 1008.1 (3.6) | 100.81 (0.36) |
| Oxycodone 1 mg/mL + S-ketamine 0.75 mg/mL | Oxycodone | 14 | 6 | 1000.0 | 1018.3 (2.4) | 101.83 (0.24) |
| Oxycodone 1 mg/mL + S-ketamine 0.75 mg/mL | Oxycodone | 21 | 6 | 1000.0 | 1018.0 (3.9) | 101.81 (0.39) |
| Oxycodone 1 mg/mL + S-ketamine 0.75 mg/mL | Oxycodone | 28 | 6 | 1000.0 | 1003.7 (4.4) | 100.37 (0.44) |
| Oxycodone 1 mg/mL + S-ketamine 0.75 mg/mL | Oxycodone | 30<sup>3</sup> | 6 | 1000.0 | 1001.8 (3.1) | 100.18 (0.31) |
| Oxycodone 1 mg/mL + S-ketamine 0.75 mg/mL | S-Ketamine | 0 | 6 | 750.0 | 750.0 (0.0) | 100.00 (0.00) |
| Oxycodone 1 mg/mL + S-ketamine 0.75 mg/mL | S-Ketamine | 3 | 6 | 750.0 | 726.9 (4.2) | 96.92 (0.56) |
| Oxycodone 1 mg/mL + S-ketamine 0.75 mg/mL | S-Ketamine | 7 | 6 | 750.0 | 740.7 (1.8) | 98.77 (0.24) |
| Oxycodone 1 mg/mL + S-ketamine 0.75 mg/mL | S-Ketamine | 14 | 6 | 750.0 | 740.3 (3.2) | 98.70 (0.42) |
| Oxycodone 1 mg/mL + S-ketamine 0.75 mg/mL | S-Ketamine | 21 | 6 | 750.0 | 724.6 (3.9) | 96.62 (0.52) |
| Oxycodone 1 mg/mL + S-ketamine 0.75 mg/mL | S-Ketamine | 28 | 6 | 750.0 | 721.8 (3.8) | 96.23 (0.51) |
| Oxycodone 1 mg/mL + S-ketamine 0.75 mg/mL | S-Ketamine | 30<sup>3</sup> | 6 | 750.0 | 718.5 (3.6) | 95.80 (0.48) |
| Oxycodone 1 mg/mL + dexmedetomidine 2.5 µg/mL | Oxycodone | 0 | 6 | 1000.0 | 1000.0 (0.0) | 100.00 (0.00) |
| Oxycodone 1 mg/mL + dexmedetomidine 2.5 µg/mL | Oxycodone | 3 | 6 | 1000.0 | 1001.3 (4.5) | 100.13 (0.45) |
| Oxycodone 1 mg/mL + dexmedetomidine 2.5 µg/mL | Oxycodone | 7 | 6 | 1000.0 | 1015.9 (11.5) | 101.59 (1.15) |
| Oxycodone 1 mg/mL + dexmedetomidine 2.5 µg/mL | Oxycodone | 14 | 6 | 1000.0 | 1001.2 (3.4) | 100.12 (0.34) |
| Oxycodone 1 mg/mL + dexmedetomidine 2.5 µg/mL | Oxycodone | 21 | 6 | 1000.0 | 1020.8 (7.9) | 102.08 (0.79) |
| Oxycodone 1 mg/mL + dexmedetomidine 2.5 µg/mL | Oxycodone | 28 | 6 | 1000.0 | 1010.4 (6.9) | 101.04 (0.69) |
| Oxycodone 1 mg/mL + dexmedetomidine 2.5 µg/mL | Oxycodone | 30<sup>3</sup> | 6 | 1000.0 | 1020.1 (6.3) | 102.02 (0.63) |
| Oxycodone 1 mg/mL + dexmedetomidine 2.5 µg/mL | Dexmedetomidine | 0 | 6 | 2.5 | 2.5 (0.0) | 100.00 (0.00) |
| Oxycodone 1 mg/mL + dexmedetomidine 2.5 µg/mL | Dexmedetomidine | 3 | 6 | 2.5 | 2.4 (0.0) | 97.79 (0.34) |
| Oxycodone 1 mg/mL + dexmedetomidine 2.5 µg/mL | Dexmedetomidine | 7 | 6 | 2.5 | 2.4 (0.0) | 96.69 (1.33) |
| Oxycodone 1 mg/mL + dexmedetomidine 2.5 µg/mL | Dexmedetomidine | 14 | 6 | 2.5 | 2.4 (0.0) | 97.63 (1.32) |
| Oxycodone 1 mg/mL + dexmedetomidine 2.5 µg/mL | Dexmedetomidine | 21 | 6 | 2.5 | 2.5 (0.0) | 100.59 (1.28) |
| Oxycodone 1 mg/mL + dexmedetomidine 2.5 µg/mL | Dexmedetomidine | 28 | 6 | 2.5 | 2.4 (0.0) | 97.89 (0.65) |
| Oxycodone 1 mg/mL + dexmedetomidine 2.5 µg/mL | Dexmedetomidine | 30<sup>3</sup> | 6 | 2.5 | 2.4 (0.0) | 96.51 (0.95) |
| Oxycodone 1 mg/mL + dexmedetomidine 5 µg/mL | Oxycodone | 0 | 6 | 1000.0 | 1000.0 (0.0) | 100.00 (0.00) |
| Oxycodone 1 mg/mL + dexmedetomidine 5 µg/mL | Oxycodone | 3 | 6 | 1000.0 | 1019.0 (6.4) | 101.91 (0.64) |
| Oxycodone 1 mg/mL + dexmedetomidine 5 µg/mL | Oxycodone | 7 | 6 | 1000.0 | 1043.7 (7.0) | 104.37 (0.70) |
| Oxycodone 1 mg/mL + dexmedetomidine 5 µg/mL | Oxycodone | 14 | 6 | 1000.0 | 1019.8 (7.3) | 101.98 (0.73) |
| Oxycodone 1 mg/mL + dexmedetomidine 5 µg/mL | Oxycodone | 21 | 6 | 1000.0 | 1041.0 (8.8) | 104.10 (0.88) |
| Oxycodone 1 mg/mL + dexmedetomidine 5 µg/mL | Oxycodone | 28 | 6 | 1000.0 | 1032.5 (5.1) | 103.25 (0.51) |
| Oxycodone 1 mg/mL + dexmedetomidine 5 µg/mL | Oxycodone | 30<sup>3</sup> | 6 | 1000.0 | 1040.8 (7.3) | 104.08 (0.73) |
| Oxycodone 1 mg/mL + dexmedetomidine 5 µg/mL | Dexmedetomidine | 0 | 6 | 5.0 | 5.0 (0.0) | 100.00 (0.00) |
| Oxycodone 1 mg/mL + dexmedetomidine 5 µg/mL | Dexmedetomidine | 3 | 6 | 5.0 | 5.0 (0.1) | 100.38 (1.11) |
| Oxycodone 1 mg/mL + dexmedetomidine 5 µg/mL | Dexmedetomidine | 7 | 6 | 5.0 | 5.2 (0.0) | 103.22 (0.96) |
| Oxycodone 1 mg/mL + dexmedetomidine 5 µg/mL | Dexmedetomidine | 14 | 6 | 5.0 | 5.1 (0.1) | 102.37 (1.33) |
| Oxycodone 1 mg/mL + dexmedetomidine 5 µg/mL | Dexmedetomidine | 21 | 6 | 5.0 | 5.1 (0.0) | 102.83 (0.76) |
| Oxycodone 1 mg/mL + dexmedetomidine 5 µg/mL | Dexmedetomidine | 28 | 6 | 5.0 | 5.0 (0.0) | 100.19 (0.42) |
| Oxycodone 1 mg/mL + dexmedetomidine 5 µg/mL | Dexmedetomidine | 30<sup>3</sup> | 6 | 5.0 | 4.9 (0.1) | 98.72 (1.22) |
| Oxycodone 1 mg/mL + dexmedetomidine 10 µg/mL | Oxycodone | 0 | 6 | 1000.0 | 1000.0 (0.0) | 100.00 (0.00) |
| Oxycodone 1 mg/mL + dexmedetomidine 10 µg/mL | Oxycodone | 3 | 6 | 1000.0 | 1026.9 (6.6) | 102.69 (0.66) |
| Oxycodone 1 mg/mL + dexmedetomidine 10 µg/mL | Oxycodone | 7 | 6 | 1000.0 | 1038.5 (3.9) | 103.85 (0.39) |
| Oxycodone 1 mg/mL + dexmedetomidine 10 µg/mL | Oxycodone | 14 | 6 | 1000.0 | 1025.3 (6.9) | 102.53 (0.69) |
| Oxycodone 1 mg/mL + dexmedetomidine 10 µg/mL | Oxycodone | 21 | 6 | 1000.0 | 1045.3 (5.5) | 104.53 (0.55) |
| Oxycodone 1 mg/mL + dexmedetomidine 10 µg/mL | Oxycodone | 28 | 6 | 1000.0 | 1035.8 (7.2) | 103.58 (0.72) |
| Oxycodone 1 mg/mL + dexmedetomidine 10 µg/mL | Oxycodone | 30<sup>3</sup> | 6 | 1000.0 | 1046.5 (6.6) | 104.64 (0.66) |
| Oxycodone 1 mg/mL + dexmedetomidine 10 µg/mL | Dexmedetomidine | 0 | 6 | 10.0 | 10.0 (0.0) | 100.00 (0.00) |
| Oxycodone 1 mg/mL + dexmedetomidine 10 µg/mL | Dexmedetomidine | 3 | 6 | 10.0 | 10.2 (0.1) | 102.22 (0.88) |
| Oxycodone 1 mg/mL + dexmedetomidine 10 µg/mL | Dexmedetomidine | 7 | 6 | 10.0 | 10.2 (0.1) | 101.58 (1.09) |
| Oxycodone 1 mg/mL + dexmedetomidine 10 µg/mL | Dexmedetomidine | 14 | 6 | 10.0 | 10.2 (0.1) | 102.43 (0.96) |
| Oxycodone 1 mg/mL + dexmedetomidine 10 µg/mL | Dexmedetomidine | 21 | 6 | 10.0 | 10.4 (0.1) | 104.00 (0.98) |
| Oxycodone 1 mg/mL + dexmedetomidine 10 µg/mL | Dexmedetomidine | 28 | 6 | 10.0 | 10.2 (0.2) | 101.73 (1.67) |
| Oxycodone 1 mg/mL + dexmedetomidine 10 µg/mL | Dexmedetomidine | 30<sup>3</sup> | 6 | 10.0 | 10.1 (0.1) | 101.42 (1.10) |

<sup>1</sup> Values are presented as mean (SD) of six replicate measurements.
<sup>2</sup> Concentrations are expressed as percentage of the initial concentration measured on day 0.
<sup>3</sup> Day 30 includes 28 days of refrigerated storage followed by 2 days at room temperature.

